# SUV39H1 Preserves Cancer Stem Cell Chromatin State and Properties in Glioblastoma

**DOI:** 10.1101/2024.08.15.607856

**Authors:** Chunying Li, Qiqi Xie, Sugata Ghosh, Bihui Cao, Yuanning Du, Giau Van Vo, Timothy Y. Huang, Charles Spruck, Y. Alan Wang, Kenneth P. Nephew, Jia Shen

## Abstract

Of the more than 100 types of brain cancer, glioblastoma (GBM) is the deadliest. As GBM stem cells (GSCs) are considered to be responsible for therapeutic resistance and tumor recurrence, effective targeting and elimination of GSCs could hold promise for preventing GBM recurrence and achieving potential cures. We show here that *SUV39H1*, which encodes a histone-3, lysine-9 methyltransferase, plays a critical role in GSC maintenance and GBM progression. Upregulation of SUV39H1 was observed in GBM samples compared to normal brain tissues, and knockdown of SUV39H1 in patient-derived GSCs impaired their proliferation and stemness. Single-cell RNA-seq analysis demonstrated restricted expression of SUV39H1 is in GSCs relative to non-stem GBM cells, likely due to super-enhancer-mediated transcriptional activation, while whole cell RNA-seq analysis revealed that SUV39H1 regulates G2/M cell cycle progression, stem cell maintenance, and cell death pathways in GSCs. By integrating the RNA-seq data with ATAC-seq (assay for transposase-accessible chromatin followed by sequencing), we further demonstrated altered chromatin accessibility in key genes associated with these pathways following SUV39H1 knockdown. Treatment with chaetocin, a SUV39H1 inhibitor, mimicked the functional effects of SUV39H1 knockdown in GSCs and sensitized GSCs to the GBM chemotherapy drug temozolomide. Furthermore, targeting SUV39H1 in vivo using a patient-derived xenograft model for GBM inhibited GSC-driven tumor formation. This is the first report demonstrating a critical role for SUV39H1 in GSC maintenance. SUV39H1-mediated targeting of GSCs could enhance the efficacy of existing chemotherapy, presenting a promising strategy for improving GBM treatment and patient outcomes.

**Highlights:**

- SUV39H1 is upregulated in GBM, especially GSCs
- Targeting SUV39H1 disrupts GSC maintenance and sensitizes GSCs to TMZ
- Targeting SUV39H1 alters chromatin accessibility at cell cycle and stemness genes
- Targeting SUV39H1 suppresses GSC-driven tumors in a patient-derived xenograft model

## 1. Introduction

Glioblastoma (GBM) is the most common primary malignant brain tumor in adults, known for its aggressiveness and lethality [1]. Despite standard treatment that includes a combination of surgery, radiotherapy, and/or temozolomide (TMZ), the prognosis for GBM patients remains poor, with a median survival of less than 16 months [2; 3]. There is an urgent need for more effective treatments to improve patient survival.

The resistance of GBM to standard treatment is partly attributed to intratumoral heterogeneity driven by GBM stem cells (GSCs), which possess self-renewal and differentiation properties, and exhibit strong tumorigenic potential [4; 5; 6]. GSCs can differentiate into various cell types within GBM, such as endothelial cells, pericytes, and non-stem GBM cells (NSGCs). As GSCs can initiate and propagate tumors, resist standard therapies including TMZ, repopulate tumors after treatment and contribute to disease relapse, eliminating GSCs could overcome chemoresistance and represent an effective therapeutic strategy in GBM.

Epigenetic modifications, including DNA methylation and histone modifications, profoundly impact gene expression and cellular behavior in normal and cancer cells, including stem cells [7; 8]. Targeting epigenetic regulators in GSCs has the potential to reverse aberrant gene expression patterns, disrupt stemness, and sensitize GSCs to existing therapies, offering new avenues for effective GBM treatment. Previous studies have identified several epigenetic regulators involved in maintaining GSC stemness and therapeutic resistance, including PRMT6 [9], PRC2 [10; 11], KDM2B [12], DNMT1 [13], HDACs [14; 15], and HELLS [16]. SUV39H1 is a key epigenetic regulator and histone methyltransferase responsible for trimethylating histone H3 at lysine 9 (H3K9me3) [17; 18], which promotes heterochromatin formation and transcriptional repression. However, the specific roles of SUV39H1 in GSCs and GBM remains to be determined.

To address this gap, we employed an integrative approach, combining in vitro and in vivo experiments with bioinformatics analyses, to investigate the functional significance of SUV39H1 in GSCs and GBM progression.

## 2. Materials and methods

### 2.1. Cell culture

Lenti-X-293T cells (Takara Bio, cat# 632180) were cultured in DMEM media (Gibco, cat# 11995065) supplemented with 10% FBS (Corning, cat# 35015CV). Patient-derived GSC3565 and GSC1914 cells were provided by Dr. Jeremy Rich (UPMC Hillman Cancer Center, Pittsburgh, PA, USA). GSC3565 cells originated from a 32-year-old male GBM patient, and GSC1914 cells from a 64-year-old male GBM patient [19]. These GSCs were cultured as previously described [20]. To minimize cell culture artifacts, patient-derived xenografts were produced and maintained as a renewable source of tumor cells.

### 2.2. Lentivirus preparation and infection

Lenti-X-293T cells cultured in DMEM media were transfected with lentivirus packaging plasmids (pMD2.g and psPAX2), along with either control or SUV39H1 shRNA vectors (Sigma-Aldrich Mission shRNA, #1 TRCN0000285355 and #2 TRCN0000275322), or pHAGE PGK-GFP-IRES-LUC-W (Addgene, cat# 46793) using the jetPRIME transfection reagent (Polyplus, cat# 101000001). After 24 hrs, the DMEM media was replaced with GSC culture media, and the cells were incubated for an additional 48 hrs at 37°C to produce lentivirus. The supernatant containing the lentivirus was harvested and saved in - 80°C. For lentivirus infection, GSCs (4 x 10^5^ cells per well in a 6-well plate) were incubated with lentivirus for 18 hrs, followed by a media change to GSC culture media and additional 48 hrs incubation.

### 2.3. Cell cycle analysis

GSC3565 or GSC1914 cells transduced with shRNAs were dissociated into single cells using accutase solution (Innovative Cell Technologies, cat# AT104), washed with cold PBS, and fixed in cold 100% ethanol and stored at −20°C for >24 hrs. After washing with cold PBS, the cells were stained with FxCycle PI/RNase Staining Solution (Invitrogen, cat# F10797) for 15 mins at room temperature, following the manufacturer’s instructions. Cells were then analyzed using a CytoFlex LX flow cytometer (Beckman Coulter). Cell cycle data were analyzed using FlowJo software (BD Biosciences).

### 2.4. Tumorsphere assay and extreme limiting dilution assay (ELDA)

For tumorsphere assays, GSCs expressing the indicated shRNAs were digested into single cells and seeded into 6-well plates (4 x 10^5^ per well for GSC3565, 1 x 10^5^ per well for GSC1914). After 72 hrs, the cells were imaged using an Axiovert 40 CFL inverted microscope (Zeiss), and tumorspheres were counted in selected fields. For chaetocin treatments, the GSCs were seeded into wells and treated with either vehicle or 50 nM chaetocin for 48 hrs before imaging. ELDA is an assay that quantifies the frequency of self-renewing cells, essential for identifying cancer stem cells, and enhances accuracy by statistically analyzing sphere-forming ability. For ELDA, GSCs transduced with shRNAs were seeded into 96-well plates at varying densities in triplicate. After 13 days of incubation, wells containing tumorspheres were counted. Stem cell frequency was estimated using the online ELDA analysis tool (http://bioinf.wehi.edu.au/software/elda/).

### 2.5. Immunohistochemistry (IHC)

Normal brain and GBM tissues (Table S1) were obtained from the Enterprise Clinical Research Operations (ECRO) Biorepository at Indiana University Health Methodist Hospital with approval from the Institutional Review Board (IRB) for the collection of human biological materials. Tissues were processed in formalin-fixed paraffin-embedded (FFPE) format, and the staining was conducted as previously described [bioRxiv2024.04.18.590173]. The antibody used was SUV39H1 (Thermo Fisher Scientific, cat# PA5-29470, 1:100). For IHC staining of brain tissues extracted from NSG mice injected with control knockdown (Ctrl KD) or SUV39H1 KD GSCs, the tissues were processed similarly to the above-described method. The antibodies used were SUV39H1 (Thermo Fisher Scientific, cat# PA5-29470, 1:100), Ki67 (Cell Signaling Technology, cat# 9449, 1:1000), and OLIG2 (EMD Millipore, cat# MABN50, 1:1000). All slides were imaged using Motic EasyScan Pro 6 scanner (Motic).

### 2.6. Western blotting

For western blotting, GSCs were lysed in cold RIPA buffer (Thermo Scientific, cat# 89901) supplemented with Protease and Phosphatase Inhibitor Mini Tablets (Thermo Scientific, cat# A32961). The procedure was detailed previously [21]. The following antibodies were used: SUV39H1 rabbit antibody (Invitrogen, cat# 702443, 1:1000), OLIG2 rabbit antibody (Cell Signaling Technology, cat# 65915, 1:1000), GFAP mouse antibody (Cell Signaling Technology, cat# 3670, 1:1000), β-Actin mouse antibody (Cell Signaling Technology, cat# 3700, 1:1000), α-Tubulin rabbit antibody (Proteintech, cat# 11224-1-AP, 1:4000), HRP-conjugated rabbit secondary antibodies (Invitrogen, cat# 31460, 1:10000), and mouse secondary antibodies (Invitrogen, cat# 31430, 1:10000).

### 2.7. Quantitative PCR (qPCR)

Total RNA was isolated using the RNeasy Plus Mini Kit (QIAGEN, cat# 74136). cDNA synthesis was performed with the qScript cDNA SuperMix (Quantabio, cat# 95048). qPCR reactions were carried out using PowerUp SYBR Green Master Mix (Applied Biosystems, cat# 25742) in CFX Duet Real-Time PCR System (Bio-Rad). The thermal cycling conditions were as follows: an initial denaturation at 95°C for 30 secs, followed by 40 cycles of 95°C for 3 secs and 60°C for 20 secs. Analyses were performed in triplicate for each data point. qPCR primers used for gene expression analysis are listed in Table S2.

### 2.8. Immunofluorescence and EdU (5-ethynyl-2’-deoxyuridine) assays

For immunofluorescence staining of GSCs, coverslips (neuVitro, cat# GG-12-laminin) were placed in 24-well plate and coated with matrigel (Corning, cat# 356231). GSCs grown on the coverslips were washed with cold PBS, fixed with 4% paraformaldehyde, permeabilized with 0.1% Triton X-100 in PBS, and incubated in blocking buffer (1% BSA in PBS). OLIG2 rabbit antibody (Cell Signaling Technology, cat# 65915, 1:200) and Alexa Fluor 488-labeled anti-rabbit secondary antibody (Invitrogen, cat# A11008, 1:500) were used. For co-staining of SUV39H1 and OLIG2 in GBM tissues, the slides were simultaneously stained with two primary antibodies: SUV39H1 (Thermo Fisher Scientific, cat# PA5-29470, 1:500) and OLIG2 (EMD Millipore, cat# MABN50, 1:1000). Alexa Fluor 488-labeled anti-rabbit secondary antibody (Invitrogen, cat# A11008, 1:500) and Alexa Fluor 594-labeled anti-mouse secondary antibody (Invitrogen, cat# A11005, 1:500) were used. For the EdU assay, GSCs grown on matrigel-coated coverslips were labeled with EdU for 4 hrs, and the subsequent steps were performed according to the instructions of the Click-iT EdU Cell Proliferation Kit (Invitrogen, cat# C10337). All images were captured using a Nikon DS-Fi3 fluorescence microscope.

### 2.9. Cell proliferation and viability assays

GSCs were seeded in 96-well plates (2 x 10^3^ cells per well), and relative cell numbers were measured on specified days using the CellTiter-Glo kit (Promega, cat# 9863) following the manufacturer’s instructions. For the in vitro drug treatment, GSC3565 cells in 96-well plates were treated with varying concentrations of TMZ (0, 200, 400, 800 nM) and chaetocin (0, 10, 20, 40 nM), either individually or in combination. After 72 hours, cell viability was measured using the CellTiter-Glo kit. For GSC1914 cells, the concentrations used were TMZ (0, 750, 1000, 1500 nM) and chaetocin (0, 20, 40, 80 nM). Synergy scores were determined using the SynergyFinder web tool (https://synergyfinder.org/). All experiments were conducted in triplicate.

### 2.10. In vivo xenograft studies

This animal study was performed under the approval of the Institutional Animal Care and Use Committee (IACUC) at Sanford Burnham Prebys Medical Discovery Institute. NSG mice (male and female, aged 4 to 6 weeks) were intracranially injected with either Ctrl KD or SUV39H1 KD GSC3565-luc cells (expressing luciferase; 5 x 10^4^ cells per mouse). Tumor size was monitored via luciferase signal using a IVIS Spectrum Imager (Xenogen). Animals were sacrificed if weight loss exceeded 20% and/or they displayed neurological signs, including but not limited to lethargy, ataxia, or seizures. On day 29, the brains were extracted, and hematoxylin and eosin (H&E) staining was performed on sections for histological analysis. IHC staining of SUV39H1, OLIG2, and KI-67 was also performed.

### 2.11. RNA-seq

Total RNA isolated from GSC3565 and GSC1914 cells with Ctrl KD or SUV39H1 KD was subjected to RNA-seq at Novogene Corporation. Quality control of the raw RNA-seq reads was performed using FastQC and fastp, and adapter sequences were removed with TrimGalore. Trimmed reads were aligned to the human reference genome (hg19) using STAR aligner. Quantification of transcript abundance was performed using Salmon in quasi-mapping mode and the resulting quantification files were imported into R for differential expression analysis using the DESeq2 package. Differentially expressed genes (DEGs) were identified with a cutoff of log_2_ fold change > 1 or < −1 and an adjusted P value < 0.05. DEGs were visualized using volcano plots generated with the ggplot2 package. Gene set enrichment analysis (GSEA) was performed using the preranked list of DEGs based on the log_2_ fold change and adjusted P value. The GSEA visualization and enrichment were conducted using the R package clusterProfiler. Enriched pathways and gene ontology (GO) terms were represented as bubble plots using Cytoscape.

### 2.12. ATAC-seq (assay for transposase-accessible chromatin followed by sequencing)

GSC3565 cells treated with vehicle or chaetocin 50 nM for 48 hrs were subjected to ATAC-seq at the Center for Medical Genomics at Indiana University School of Medicine. ATAC-seq libraries were prepared following the Omni-ATAC protocol. Briefly, GSCs were washed with cold PBS and lysed in ATAC lysis buffer (10 mM Tris-HCl, pH 7.4, 10 mM NaCl, 3 mM MgCl_2_, 0.1% Igepal CA-630). Nuclei were collected by centrifugation and resuspended in transposition reaction mix containing Tn5 transposase (Illumina, cat# FC-121-1030) and incubated at 37°C for 30 mins. The transposed DNA was purified using the DNA Clean & Concentrator-5 (Zymo, cat# D4003) and PCR was amplified using Nextera PCR primers. The amplified libraries were purified with the KAPA HiFi HotStart ReadyMix (Roche, cat#KK2602) and quantified using a Qubit fluorometer (Thermo Fisher Scientific). Libraries were sequenced on an Illumina platform to generate paired-end reads. Sequencing reads were trimmed for adapter sequences using Trim Galore and aligned to the human reference genome (hg19) using Bowtie2. Aligned reads were filtered to remove duplicate reads, mitochondrial reads, and low-quality reads using SAMtools and sambamba. Peaks were called using MACS2 to identify regions of open chromatin. The resulting peak files were further processed to generate a high-confidence set of peaks by filtering out blacklist regions and peaks overlapping with ENCODE blacklisted regions. The number of reads within each peak was quantified using featureCounts from the Subread package. Differentially accessible regions between conditions were identified using DESeq2, with an adjusted P value < 0.05 considered significant. Peaks were annotated to the nearest genes using HOMER, and GO enrichment analysis was performed on the annotated genes using the clusterProfiler package. Heatmaps and average signal profiles of ATAC-seq data were generated using DeepTools. The bigWig files for visualization were created using bedGraphToBigWig. The Integrative Genomics Viewer (IGV) was used to visualize the ATAC-seq signal tracks across the genome.

To integrate ATAC-seq data with RNA-seq data, the overlap of differentially accessible regions with DEGs was analyzed using BEDTools intersect. The genomic regions enrichment of annotations tool (GREAT) was used to identify the biological relevance of the genomic regions identified.

### 2.13. Single-cell RNA-seq Analysis

Single-cell RNA-seq data were downloaded from the CELLxGENE database and integrated and analyzed using Seurat (v4.4.0). Prior to integration, quality control was performed to remove low-quality cells (< 200 genes detected, > 10% mitochondrial reads) and potential doublets. Following quality control, cells were annotated based on the cell type information provided in the CELLxGENE database. Data integration was achieved using Seurat’s SCTransform workflow, followed by batch effect correction with the Harmony algorithm. Dimensionality reduction was performed using Principal Component Analysis (PCA) on the top 2000 variable genes, with the first 30 principal components used for downstream analysis. Cells were clustered based on their gene expression profiles using the Louvain algorithm with a resolution of 0.8. Uniform Manifold Approximation and Projection (UMAP) was used for visualization of the high-dimensional data in two dimensions. Cell type annotations were refined using canonical markers, with a comprehensive list of markers used for annotation provided in Table S3. Differential expression analysis was performed using the Wilcoxon rank-sum test, and significant genes were identified based on an adjusted P value < 0.05 and log_2_ fold change > 1, with Benjamini-Hochberg correction for multiple testing. Specific markers for GSCs (including OLIG2, SOX2, NES) and other cell types were visualized on UMAP plots using a color gradient to represent expression intensity.

### 2.14. Statistical analysis

All statistical analyses were conducted using GraphPad Prism software, with a P value of less than 0.05 considered statistically significant. Data are presented as mean ± SD. The statistical methods for each experiment are detailed in the figure legends.

## 3. Results

### 3.1. SUV39H1 is upregulated in GBM

We first investigated the expression of SUV39H1 in GBM using RNA-seq data from GBM datasets (The Cancer Genome Atlas (TCGA), Murat, Kamoun, and Rembrandt). Compared to non-tumor brain tissues, SUV39H1 expression was significantly increased in GBM tissues (P < 0.05; Fig. 1A-D). Overexpression of SUV39H1 in GBM tissues was confirmed by IHC (P < 0.001; Fig. 1E). These findings demonstrate that SUV39H1 is upregulated in GBM compared to normal brain tissues.

**Fig. 1.**
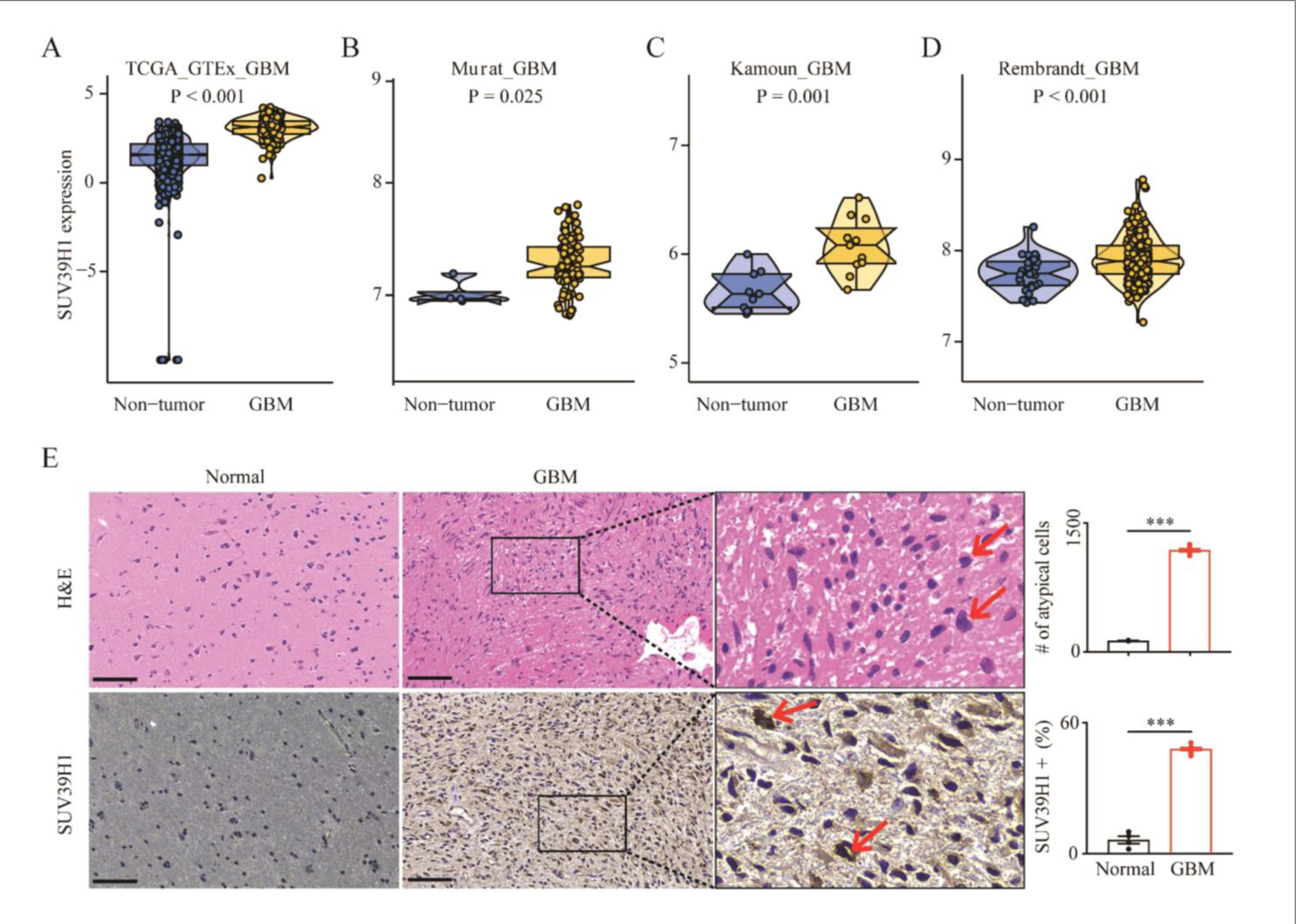
SUV39H1 is upregulated in GBM. (A-D) Violin plots showing SUV39H1 expression in non-tumor and GBM tissues across four datasets: TCGA_GTEx_GBM (A), Murat_GBM (B), Kamoun_GBM (C), and Rembrandt_GBM (D). P values were calculated using t-tests. (E) H&E staining (top row) and IHC staining of SUV39H1 (bottom row) in normal (n=4) and GBM (n=9) tissues. Quantification of the number of atypical cells and the percentage of SUV39H1-positive cells are shown on the right. Scale bars represent 100 μm. ***P < 0.001.

### 3.2. SUV39H1 is preferentially expressed in GBM stem cells

To examine the expression pattern of SUV39H1 in GBM, single-cell RNA-seq analysis was performed. UMAP clustering identified distinct cell types in GBM (Fig. 2A; Fig. S1A). We used stemness markers, including OLIG2, NES, and SOX2, to further analyze the malignant cells and distinguish GSCs from non-stem GBM cells (NSGCs) (Fig. 2B). The data revealed that SUV39H1 was preferentially overexpressed in GSCs (Fig. 2C, D). Immunofluorescence staining confirmed co-localization of SUV39H1 and OLIG2 in GBM patient tissues (Fig. 2E).

**Fig. 2.**
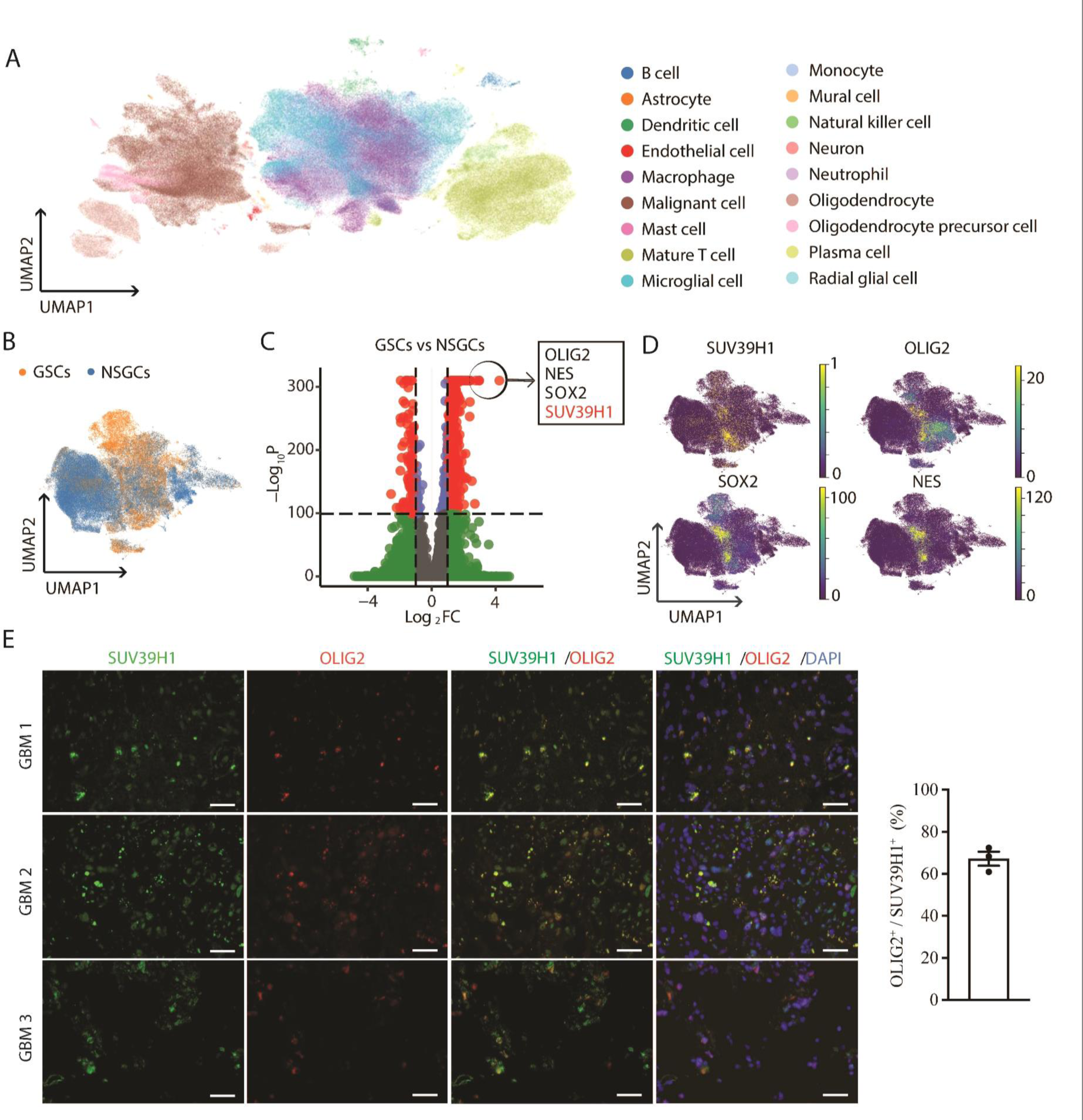
Expression patterns of SUV39H1 in GBM. (A) The UMAP clustering of single-cell RNA-seq data from GBM tumors sourced from the CELLxGENE database reveals various cell types. (B) UMAP plot distinguishing GSCs (orange) from non-stem GBM cells (NSGCs, blue). (C) Volcano plot displaying DEGs between GSCs and NSGCs. Genes with significant expression changes (|log_2_ fold change| > 2 and adjusted P value < 0.05) are shown as red dots, with the top genes (*OLIG2*, *NES*, *SOX2*, and *SUV39H1*) annotated in the box. (D) UMAP plots showing the expression patterns of the indicated genes in GSCs and NSGCs. The color intensity represents the normalized expression level. (E) Representative images (left panel) and quantification (right panel) of immunofluorescence staining showing co-localization of SUV39H1 (green) and OLIG2 (red) in GBM tissues (n=3). Scale bars represent 50 μm.

Analysis of RNA-seq data for three pairs of GSCs and their matched differentiated NSGCs (GSE54791) revealed higher SUV39H1 expression in GSCs (P < 0.01; Fig. 3A). Using GSC3565 and GSC1914 models for differentiated and undifferentiated cell states (Fig. 3B), we found that SUV39H1 levels decreased during serum-induced differentiation, accompanied by downregulation of OLIG2 and upregulation of GFAP, a differentiated NSGC marker (qPCR, Fig. 3C; western blot, Fig. 3D). ChIP-seq data of H3K27ac, an active super-enhancer mark, revealed enriched peaks at the *SUV39H1* gene in GSCs compared to differentiated NSGCs, indicating enhanced regulatory activity of SUV39H1 expression in GSCs (Fig. 3E). Additionally, RNA-seq analysis of 44 GSC models and 9 normal brain cell lines (GSE119834) demonstrated preferential SUV39H1 expression in GSCs (P = 0.004; Fig. 3F). The ChIP-seq signal tracks for H3K27ac, a histone modification linked to active enhancers that boost gene transcription, revealed increased enhancer regulation of SUV39H1 in GSCs compared to normal brain cells (Fig. 3G).

**Fig. 3.**
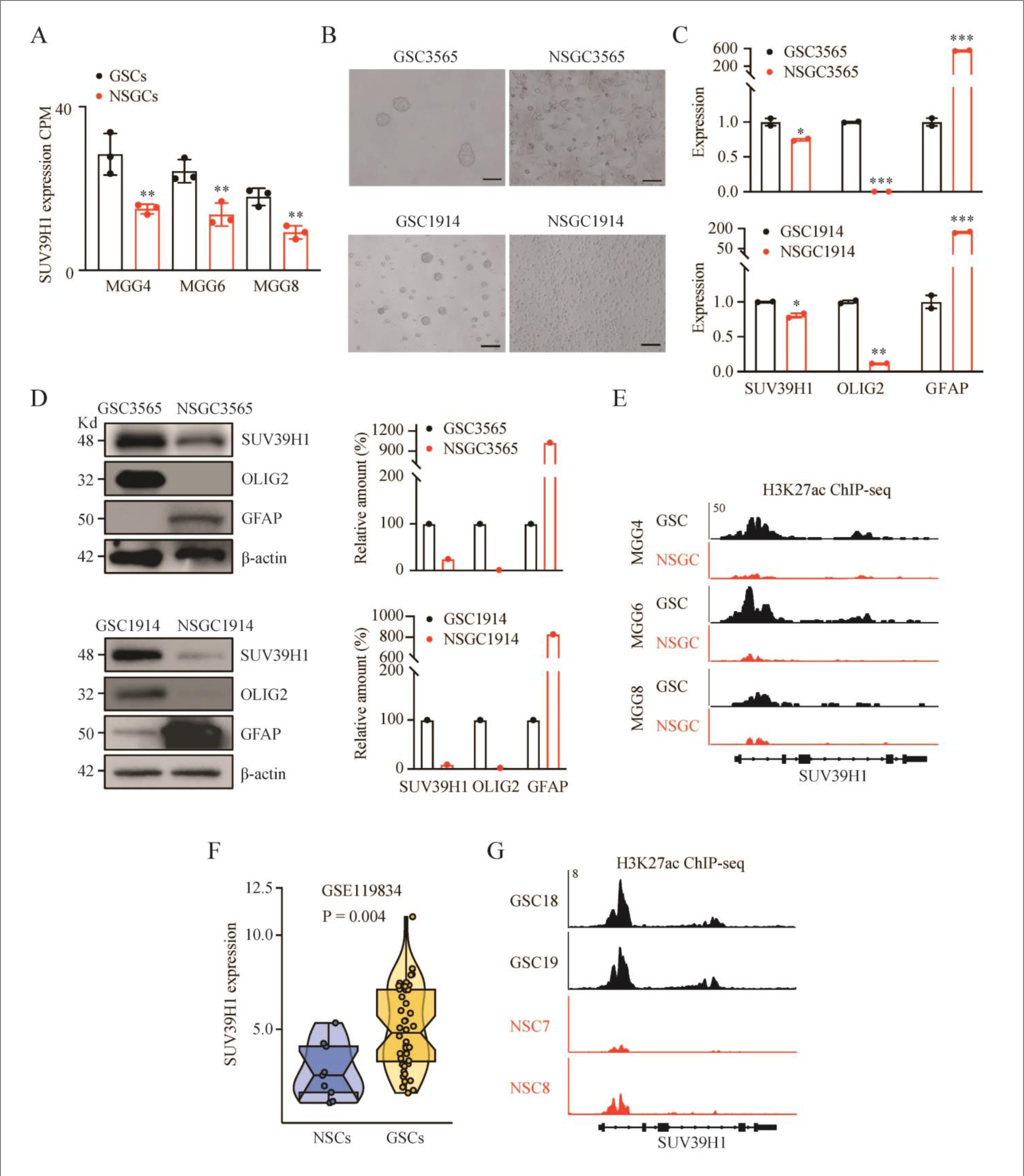
SUV39H1 expression in GSCs, NSGCs, and normal brain cells. (A) Analysis of RNA-seq data (GSE54791) for SUV39H1 expression in multiple GSC and NSGC pairs (MCG4, MCG6, MCG8). CPM: counts per million. (B-D) Representative images (B), qPCR data (C), and western blotting data and quantification (D) for GSC3565 and GSC1914 differentiation by serum induction. OLIG2 is a GSC marker, and GFAP is a differentiated NSGC marker. Scale bars represent 100 μm. (E) H3K27ac ChIP-seq data showing the *SUV39H1* locus in indicated GSCs and NSGCs. (F) Violin plot showing SUV39H1 expression levels in GSCs compared to normal stem cells (NSCs) in brain. P value was calculated using a t-test. (G) H3K27ac ChIP-seq signal tracks showing the *SUV39H1* locus in indicated GSCs and NSCs. *P < 0.05, **P < 0.01, ***P < 0.001.

### 3.3. SUV39H1 is required for GSC maintenance

SUV39H1 upregulation in GSCs suggests a potential dependency on this enzyme. To investigate the functional roles of SUV39H1 in GSCs, we first knocked down (KD) SUV39H1 in GSC3565 and GSC1914 cells using two different shRNAs and confirmed KD efficiency by western blotting (Fig. 4A). SUV39H1 KD led to decreased GSC proliferation (Fig. 4B), which was further validated by an EdU incorporation assay showing slower DNA replication in GSCs with SUV39H1 KD (Fig. 4C, D). Additionally, SUV39H1 KD impaired the self-renewal ability of GSCs. Tumorsphere formation assay showed a significant reduction in the number of tumorspheres formed by SUV39H1 KD cells (P < 0.05; Fig. 4E, F). ELDA quantitatively demonstrated a decreased stem cell frequency in SUV39H1 KD cells (P < 0.05; Fig. 4G).

**Fig. 4.**
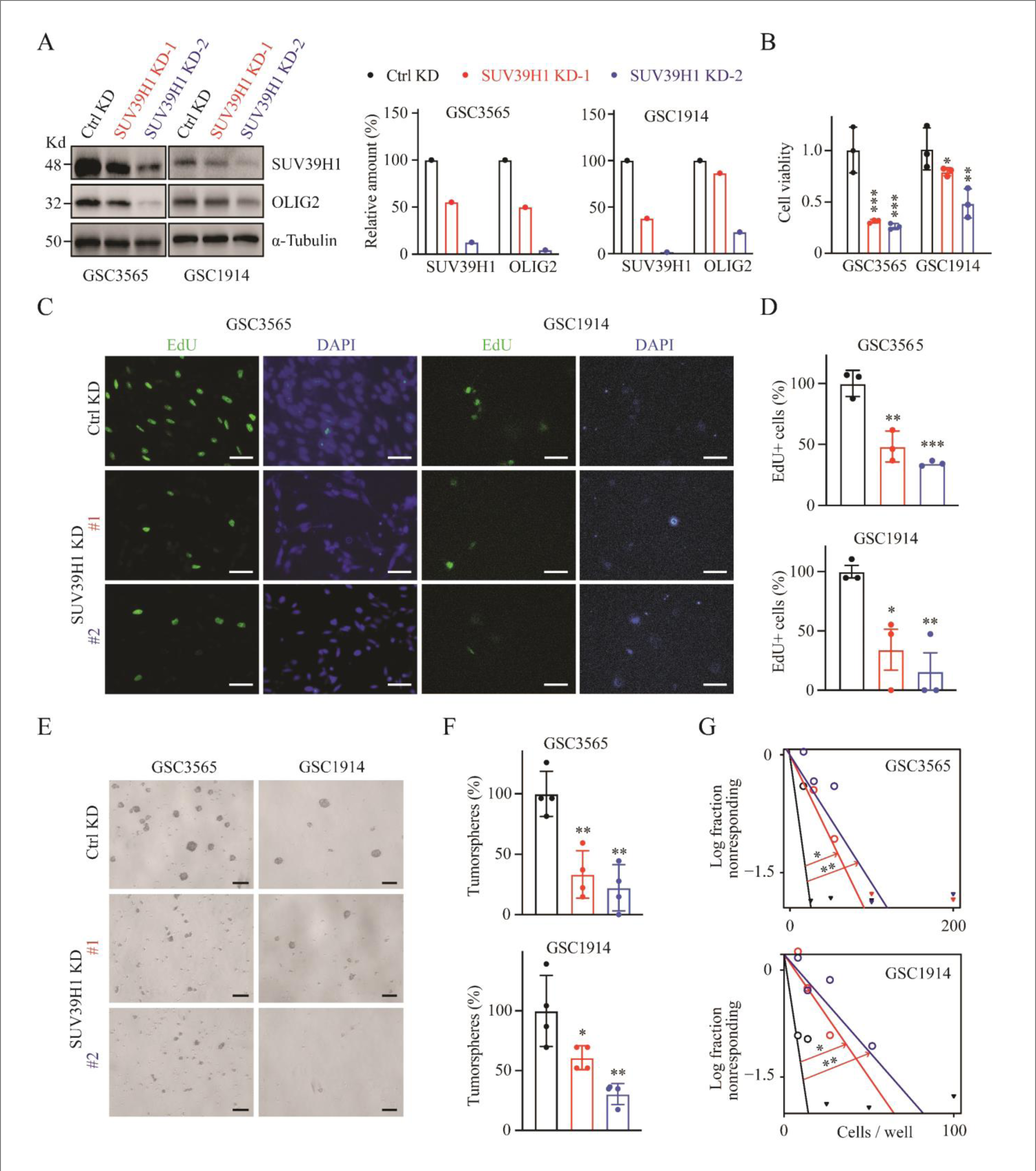
SUV39H1 regulates GSC proliferation and stemness. (A) Western blot data (left panel) and quantification (right panel) showing the levels of SUV39H1 and OLIG2 in Ctrl and SUV39H1 KD GSCs. (B) Cell viability analysis for GSC3565 (day 5) and GSC1914 (day 8) following gene knockdown. (C, D) Representative images (C) and quantification (D) of EdU incorporation assay in GSCs. Scale bars represent 50 μm. (E, F) Representative images (E) and quantification (F) of tumorsphere formation in GSCs after 72 hrs of gene knockdown. Scale bars represent 100 μm. (G) Limiting dilution assay demonstrating the self-renewal capacity of GSCs with various cell numbers after 13 days of gene knockdown. Data points represent the log fraction of wells without spheres plotted against the number of cells plated per well. *P < 0.05, **P < 0.01, ***P < 0.001, ***P < 0.001.

To further examine the effect of inhibiting SUV39H1, GSCs were treated with chaetocin, an inhibitor of SUV39H1 [22; 23]. qPCR analysis demonstrated decreased expression of SUV39H1 and OLIG2 in GSCs treated with chaetocin compared to untreated controls (Fig. 5A). Tumorsphere formation assay revealed a significant reduction in the number of tumorspheres formed in chaetocin-treated GSCs (P < 0.05; Fig. 5B, C). Notably, chaetocin treatment sensitized both GSC3565 and GSC1914 cells to the GBM chemotherapy drug TMZ, with synergy scores of 10.148 and 16.086, respectively (Fig. 5D, E). These findings demonstrate that SUV39H1 is essential for GSC maintenance.

**Fig. 5.**
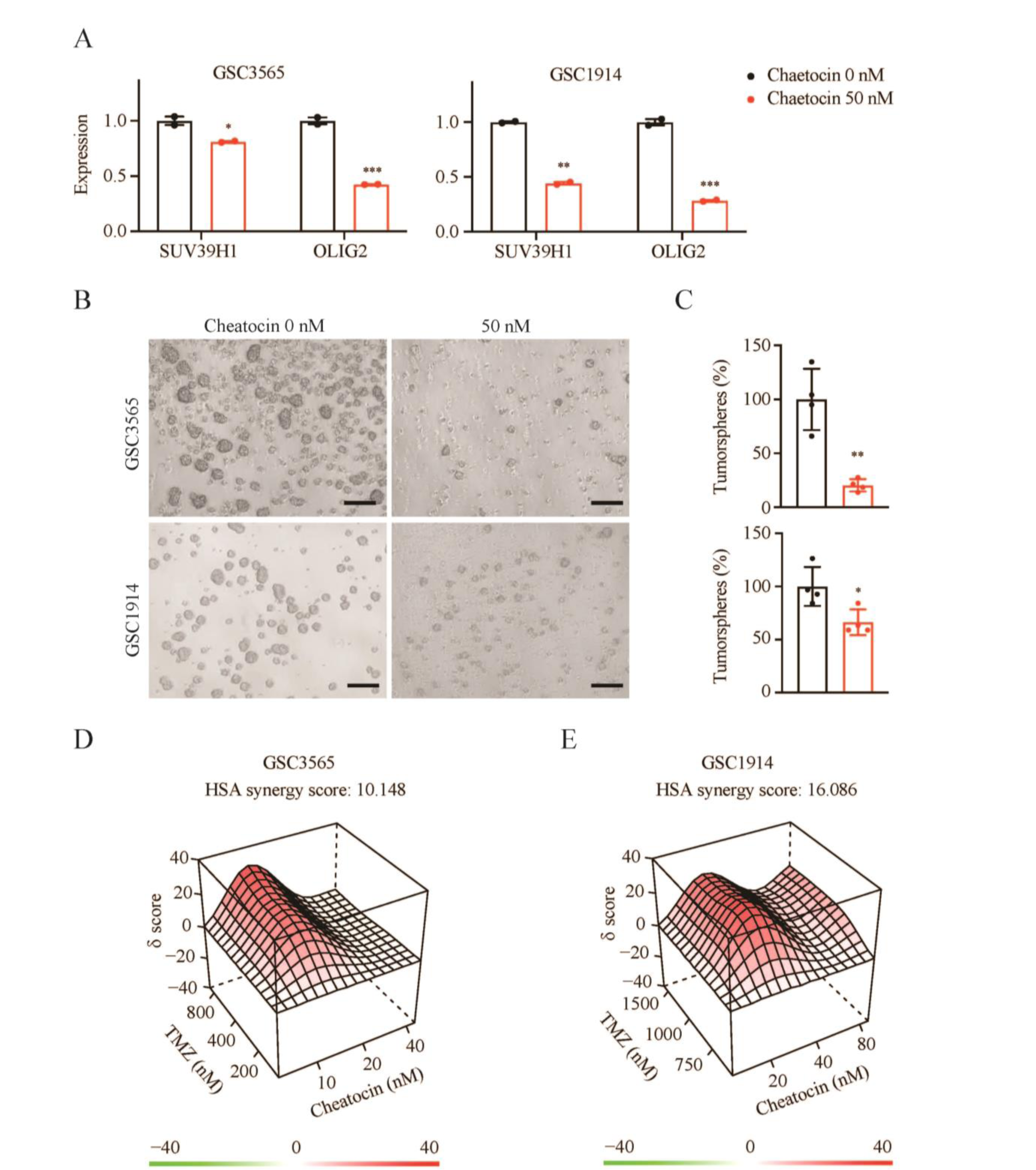
Chaetocin treatment disrupts GSCs and synergizes with TMZ. (A) qPCR analysis of SUV39H1 and OLIG2 expression in GSCs treated with 0 nM or 50 nM chaetocin. (B, C) Representative images (B) and quantification (C) of tumorsphere formation in GSCs after 48 hrs of chaetocin treatment. Scale bars represent 100 μm. (D, E) 3D synergy score plots showing the synergistic effect of chaetocin and TMZ on killing GSC3565 (D) and GSC1914 (E). *P < 0.05, **P < 0.01, ***P < 0.001.

### 3.4. SUV39H1 regulates cell cycle, stemness, and cell death pathways in GSCs

To explore the molecular mechanisms by which SUV39H1 maintains GSCs, we performed RNA-seq analysis on GSC3565 and GSC1914 cells with SUV39H1 KD versus Ctrl KD (Fig. 6A). GSEA identified several enriched pathways associated with the differentially expressed genes (DEGs) (Fig. 6B). SUV39H1 KD GSCs exhibited downregulation of G2/M cell cycle pathways (Fig. 6C). Consistently, qPCR detection of G2/M cell cycle-related genes, including *CDK16*, *CDC27*, and *CUL3*, showed decreased expression upon SUV39H1 KD (Fig. 6D). Flow cytometry analysis revealed an enrichment of the G2/M phase cell population in SUV39H1 KD GSCs (Fig. 6E). GSEA also identified decreased stem cell-related pathways in SUV39H1 KD cells (Fig. 6F), which was validated by qPCR showing downregulation of stemness genes, including *OLIG2*, *NES*, and *MYC* (Fig. 6G), and by immunofluorescence for OLIG2 (Fig. 6H). Additionally, cell death pathways were upregulated, with GSEA indicating enrichment of autophagy, ferroptosis, and pyroptosis pathways (Fig. S2A), which were validated by qPCR for selected genes (Fig. S2B). These results demonstrate that SUV39H1 regulates cell cycle, stemness, and cell death pathways in GSCs.

**Fig. 6.**
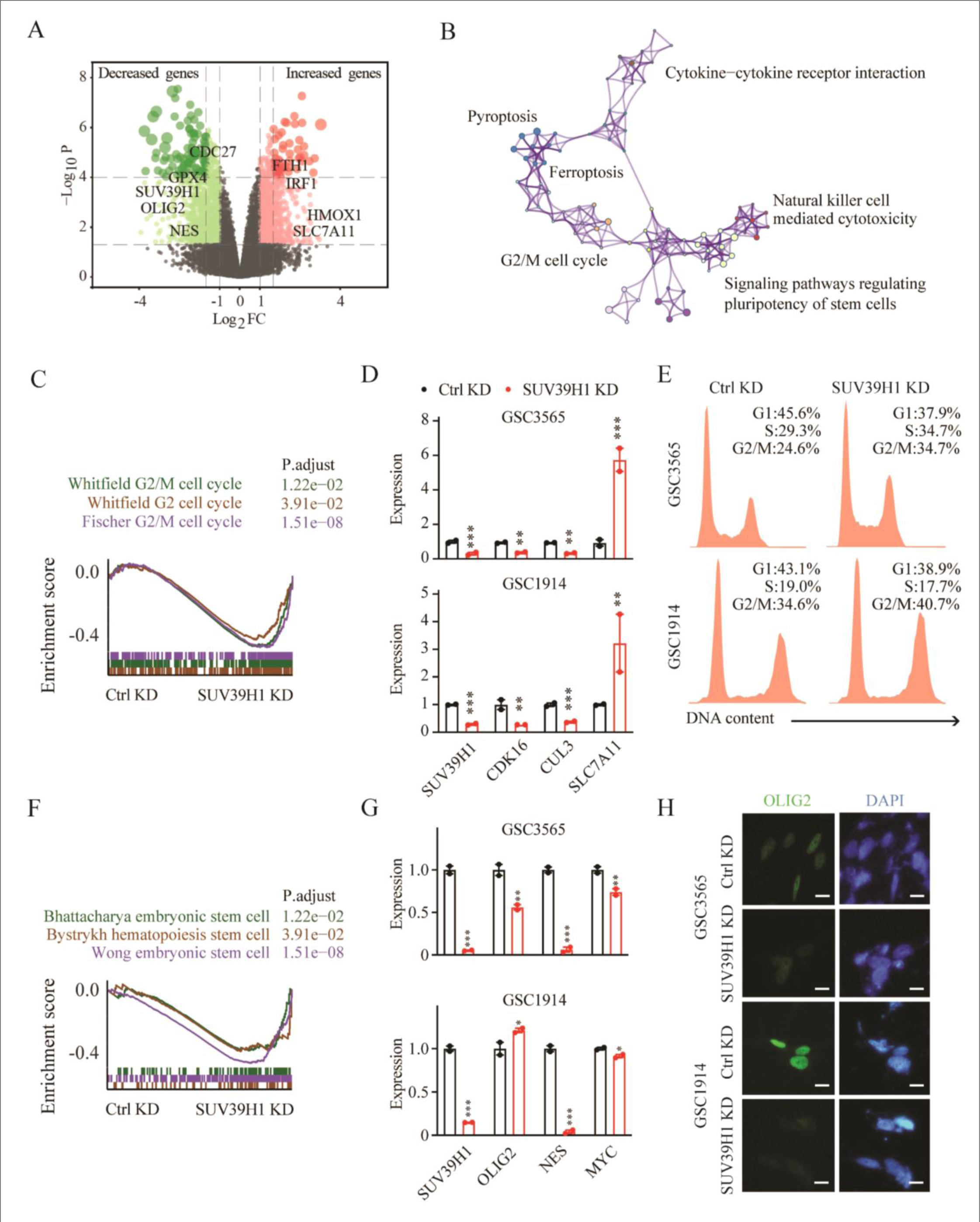
Signal pathways regulated by SUV39H1 in GSCs. (A) Volcano plot illustrating DEGs in SUV39H1 KD versus Ctrl KD GSC3565 and 1914 cells, with key genes highlighted. (B) Enrichment map visualizing the significant pathways affected by SUV39H1 KD. (C) GSEA plot showing enrichment of the G2/M cell cycle-related pathways in GSCs with SUV39H1 KD. P.adjust values indicate the significance of enrichment. (D) qPCR detection of G2/M cell cycle-related genes in GSC3565 and GSC1914 cells. (E) Flow cytometry data showing cell cycle alteration in GSCs with SUV39H1 KD. (F) GSEA plot showing enrichment of the stem cell-related pathways in SUV39H1 KD GSCs. (G) qPCR detection of stem cell-related genes in GSCs. (H) Immunofluorescence staining for OLIG2 (green) and DAPI (blue) in GSC3565 and GSC1914 cells. Scale bars represent 50 μm. *P < 0.05, **P < 0.01, ***P < 0.001.

### 3.5. Targeting SUV39H1 alters chromatin accessibility

To investigate how SUV39H1 targeting affects gene expression, we performed ATAC-seq, revealing distinct chromatin accessibility profiles between untreated and chaetocin-treated GSCs (Fig. 7A, B). These differentially accessible chromatin regions were distributed across various genomic features (Fig. 7C), and heatmap visualization highlighted a significant population of genes associated with these regions, which we named ATAC-seq-DGs (ATAC-seq-differential genes) (Fig. 7D). Integrating ATAC-seq-DGs with RNA-seq DEGs (see Fig. 6A) identified overlapping genes (n=2823, 8.4%) (Fig. 7E), with many linked to G2/M cell cycle and stem cell pathways (Fig. 7F). For example, For example, *CDC27* and *CDC6* G2/M cell cycle (Fig. 7G) and *OLIG2* and *NES* stemness genes (Fig. 7H) showed decreased chromatin accessibility and gene expression upon SUV39H1 targeting, as visualized in the ATAC-seq and RNA-seq signal tracks. These data suggest that targeting SUV39H1 reduces chromatin accessibility at these genes, resulting in downregulation of their expression, contributing to G2/M cell cycle arrest and disruption of stemness.

**Fig. 7.**
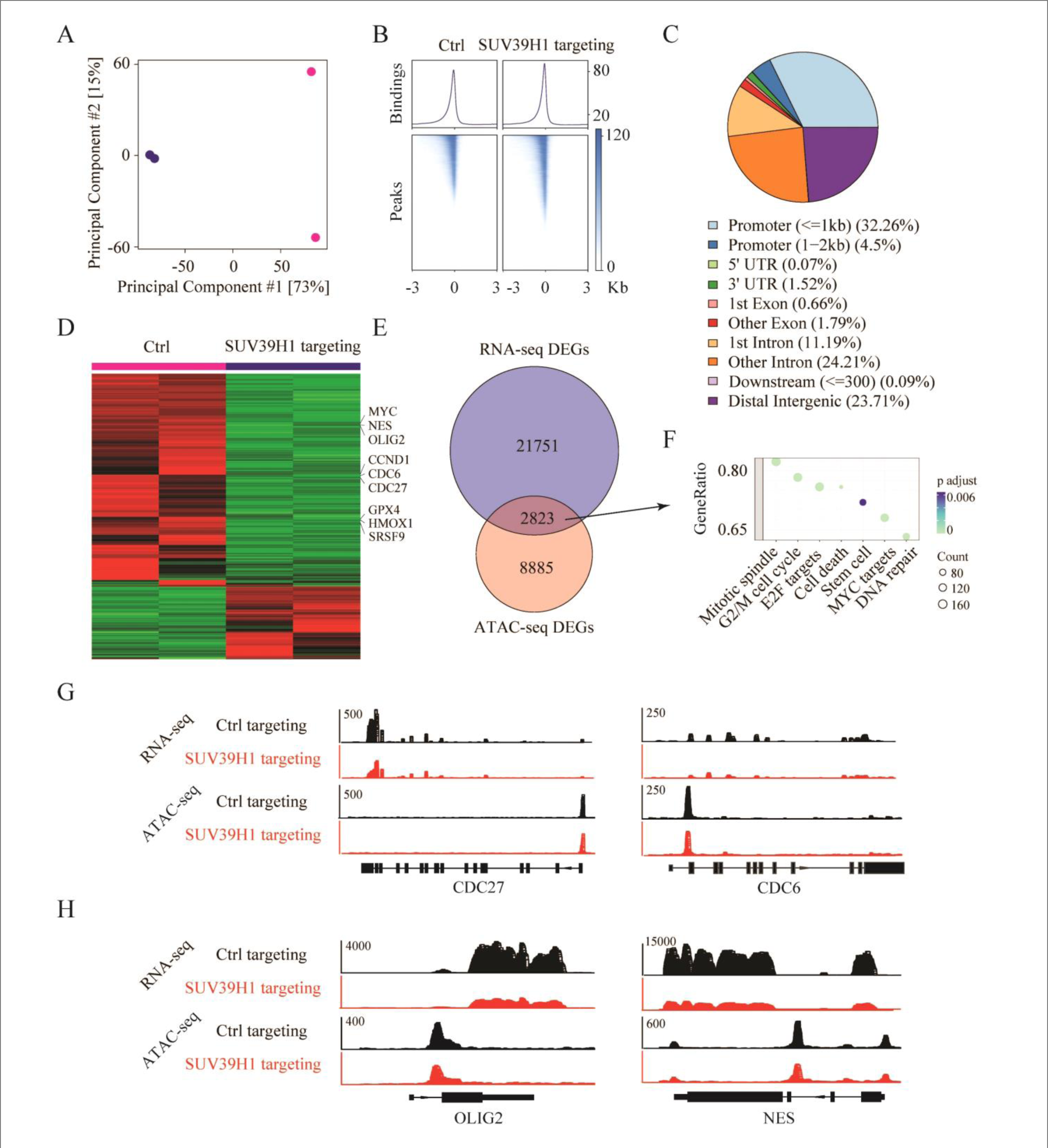
Targeting SUV39H1 alters chromatin accessibility in GSCs. (A) Principal component analysis (PCA) plot displaying the clustering of untreated (blue) and chaetocin-treated (pink) GSCs. (B) Aggregate plots (top) and heatmaps (bottom) of ATAC-seq signals at transcription start sites (TSS) in untreated and chaetocin-treated GSCs. (C) Pie chart illustrating the distribution of differentially accessible chromatin regions across various genomic features in response to chaetocin treatment. (D) Heatmap demonstrating the genes associated with altered chromatin accessibility upon SUV39H1 targeting. (E) Venn diagram showing the overlap of genes (n=2823, 8.4%) between RNA-seq DEGs (blue) and genes associated with ATAC-seq differentially accessible regions (pink) after chaetocin treatment. (F) Bubble plot of GO enrichment analysis for the overlapping genes identified in (E). (G, H) UCSC genome browser tracks displaying RNA-seq and ATAC-seq signals for indicated genes related to G2/M cell cycle (G) and stem cell maintenance (H).

### 3.6. Targeting SUV39H1 decreases GSC-driven GBM growth in mice

The impact of targeting SUV39H1 on the in vivo tumor formation ability of GSCs was assessed using a xenograft mouse model. Ctrl or SUV39H1 KD GSC3565 cells expressing luciferase were intracranially injected into the brains of immunodeficient mice (Fig. 8A, B). SUV39H1 KD GSC-derived tumors displayed significantly reduced growth in mice on day 29 (Fig. 8C, D). IHC analysis revealed decreased staining for SUV39H1, Ki67, and OLIG2 in SUV39H1 KD tumors compared to the controls (P < 0.001; Fig. 8E, F), demonstrating that targeting SUV39H1 can disrupt GSC proliferation and stemness, thereby inhibiting GBM growth in vivo.

**Fig. 8.**
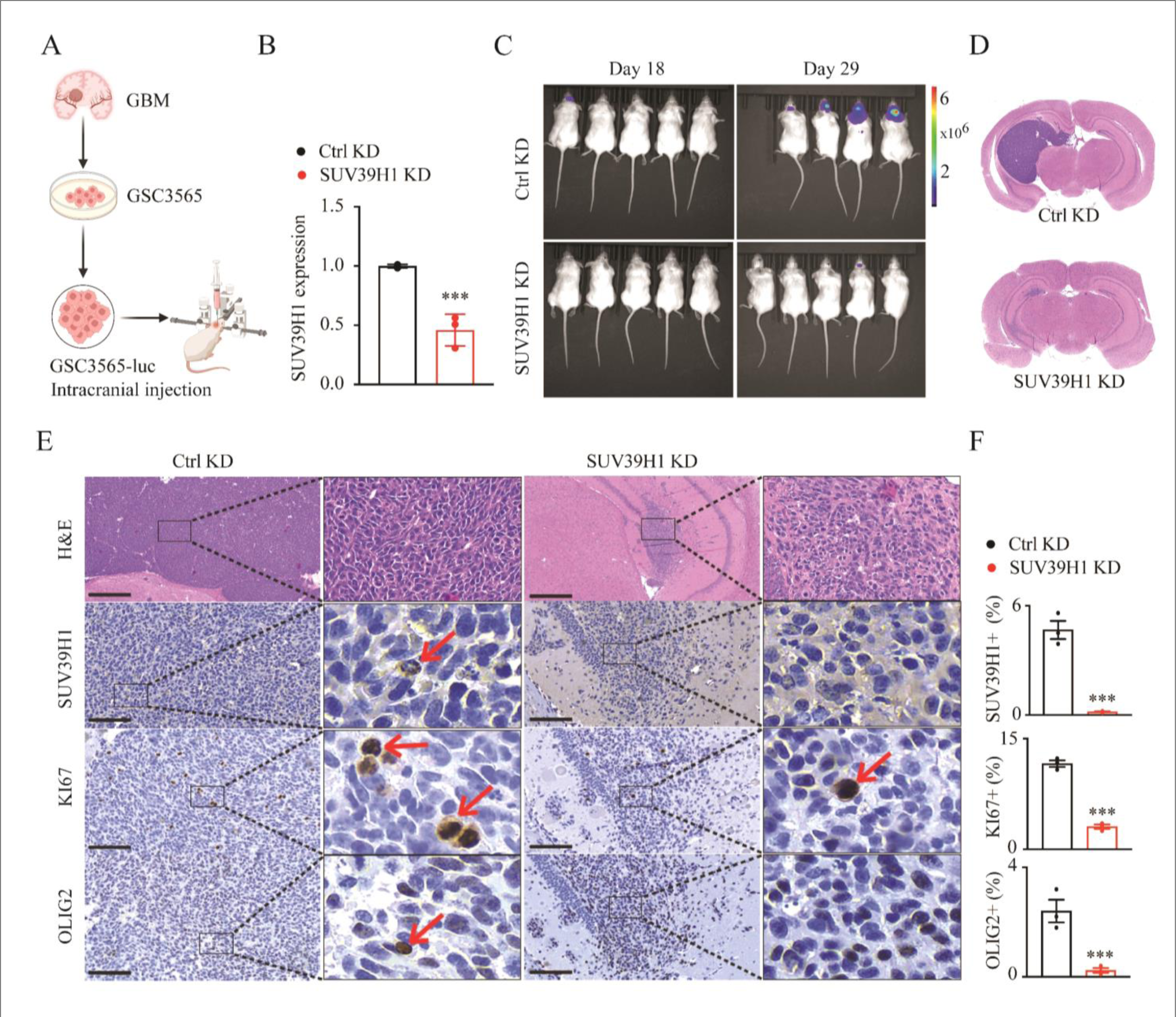
Targeting SUV39H1 decreases GSC-driven GBM growth in mice. (A) Schematic representation of the experimental design for intracranial injection of GSCs with Ctrl or SUV39H1 KD. (B) qPCR analysis of SUV39H1 expression in GSCs prior to injection. (C) Bioluminescence imaging of tumor-bearing mice on days 18 and 29 post-implantation of GSCs. (D) Representative H&E-stained coronal brain sections of mice implanted with indicated GSCs on day 29. (E, F) Representative images (E) and quantification (F) of IHC staining for SUV39H1, Ki-67, and OLIG2 in tumor sections from (D). Scale bars represent 100 μm. ***P < 0.001.

## 4. Discussion

GSCs are pivotal in GBM tumorigenesis and recurrence. GSCs exhibit robust resistance to standard-of-care treatments, including chemotherapy and radiation, largely due to their enhanced DNA repair capabilities [24], active drug efflux pumps [25; 26], and quiescent state that renders them less susceptible to treatments targeting rapidly dividing cells [27; 28]. To overcome these challenges, numerous studies have focused on targeting the genes and signaling pathways that regulate stemness and survival, with the intent to directly kill GSCs or sensitize them to conventional therapies. This study identifies SUV39H1 as a critical dependency in GSCs, regulating key pathways involved in cell cycle progression, stem cell maintenance, and cell death. Targeting SUV39H1 leads to reduced GSC maintenance, increased sensitivity to TMZ chemotherapy, and impaired GBM tumor growth in mice (Fig. 9).

**Fig. 9.**
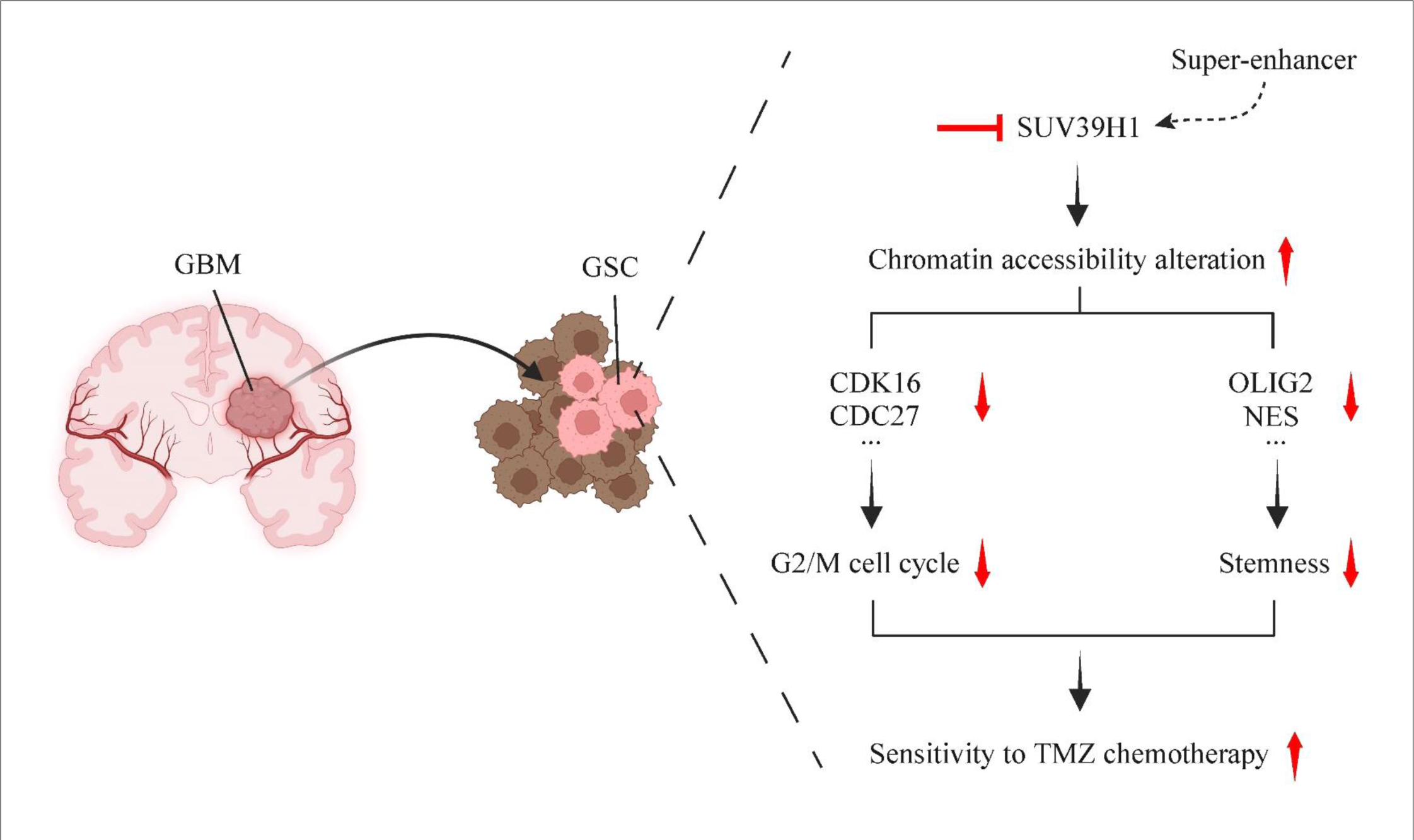
Schematic model of SUV39H1 targeting in GSCs. This working model illustrates the signaling cascade involving the super-enhancer, SUV39H1, chromatin accessibility, G2/M cell cycle regulation, stemness, and sensitivity to TMZ chemotherapy.

SUV39H1 plays a crucial role in the development and progression of various cancers. For instance, SUV39H1-mediated H3K9me3 can silence the tumor suppressor gene *p16INK4a*, promoting uncontrolled cell proliferation in acute myeloid leukemia (AML) and lung cancer cells [29; 30]. SUV39H1 deposits H3K9me3 at the promoter of the cell death receptor *FAS* gene, repressing expression and inhibiting apoptosis in metastatic colorectal cancer cells [31]. Our previous study revealed that SUV39H1 signaling contributes to the maintenance of repetitive elements and genomic stability across various cancer cell types [21]. In this study, we investigated the functional significance of SUV39H1 in cancer stem cells in GBM. We found that, in addition to regulating cell proliferation, SUV39H1 is essential for GSC stemness maintenance through regulation of chromatin accessibility. Interestingly, a previous study in bladder cancer stem cells showed that SUV39H1 maintains the self-renewal of these cells by repressing the transcription factor GATA3, leading to the upregulation of STAT3 and maintaining stemness [32]. More recently, another study linked SUV39H1 with cancer stemness in diffuse intrinsic pontine gliomas, reporting that targeting SUV39H1 led to the downregulation of growth factor receptor signaling and stemness-related programs, resulting in inhibited glioma cell growth and increased cell death [22]. However, whether SUV39H1 also regulates stemness maintenance in cancer stem cells of other cancer types remains to be determined in future studies.

In this study, we utilized single-cell RNA-seq to map the transcriptomic landscape of GBM, identifying distinct cell populations and revealing the expression of SUV39H1 across various cell types. While the primary focus was on the role of SUV39H1 in GSCs, it is evident that SUV39H1 also regulates the functions of other cell types in the GBM microenvironment. Previous research showed in tumor-infiltrating cytotoxic T lymphocytes (CTLs) in human colon carcinoma that SUV39H1 mediated H3K9me3 marks at the promoters of CTL effector genes (e.g., *GZMB*, *PRF1*, *FASLG*, and *IFNG*). This enrichment repressed their expression and diminished CTL cytotoxicity, thereby facilitating tumor immune evasion. Targeting SUV39H1 increased the expression of these effectors in CTLs, resulting in suppression of colon tumor growth [33]. Another study in diffuse large B-cell lymphoma revealed that SUV39H1 upregulated CD86+ and CD163+ tumor-associated macrophages, suggesting a novel mechanism for lymphoma progression [34]. Recent studies also underscore the role of SUV39H1 in regulating chimeric antigen receptor (CAR) T cells. Targeting SUV39H1 enhanced the persistence and antitumor effects of CAR T cells in lung and disseminated solid tumor models [35], as well as in leukemia and prostate cancer models [36]. Understanding the roles of SUV39H1 in various cell types within the tumor microenvironment is crucial for evaluating the therapeutic potential of SUV39H1 inhibitors for GBM treatment. Ideally, these inhibitors should selectively target cancer cells and pro-tumor cells, such as regulatory T (Treg) cells and myeloid-derived suppressor cells (MDSCs), while sparing or even benefiting anti-tumor cells, such as CTLs and natural killer (NK) cells.

In conclusion, our integrative approach, combining in vitro and in vivo experiments with bioinformatics analyses, provides compelling evidence that SUV39H1 is a key regulator of GSC functions. Targeting SUV39H1 could offer insights into novel strategies to improve outcomes for GBM patients.

## Data availability

The RNA-seq data were deposited in the NCBI’s Gene Expression Omnibus (GEO) database under accession number GSE273012. The ATAC-seq data were deposited in the GEO database under accession number GSE273013. Other data described in this article are available upon request from the corresponding authors.

## CRediT authorship contribution statement

**Chunying Li:** Investigation, Visualization, Methodology. **Qiqi Xie:** Investigation, Visualization, Data curation, writing–original draft. **Sugata Ghosh:** Investigation, Methodology, Writing–original draft. **Bihui Cao:** Investigation, Visualization, Methodology. **Yuanning Du:** Investigation, Visualization, Methodology, Writing–original draft. **Giau Vo:** Investigation. **Timothy Y. Huang:** Resources. **Charles Spruck:** Resources, Writing–review and editing. **Y. Alan Wang:** Writing–review and editing. **Kenneth P. Nephew:** Resources, Writing–review and editing. **Jia Shen:** Conceptualization, Methodology, Funding acquisition, Supervision, Project administration, Writing–original draft, Writing–review and editing.

## Acknowledgments

Research reported in this publication was partially supported by the National Cancer Institute of the National Institutes of Health under Award Number P30CA030199. The content is solely the responsibility of the authors and does not necessarily represent the official views of the National Institutes of Health. This study was supported by the Indiana University School of Medicine Start-Up Fund and the Schwarz Family and Friends Cancer Research Fund, both awarded to Jia Shen. Y. Alan Wang is supported by NIH R01 CA231349. We extend our gratitude to the Genomics Core Facility and Histology Core Facility at the Sanford Burnham Prebys Medical Discovery Institute for their research support. We also thank Christiane Hassel and the Indiana University Bloomington Flow Cytometry Facility for their assistance. The CytoFLEX LX instrument was partially funded by the Indiana University Office of Research through the Research Equipment Fund. We appreciate Dr. Jeremy Rich at the UPMC Hillman Cancer Center for generously providing the GSCs. Thanks also to Drs. Peter Hollenhorst, Heather M. O’Hagan, Richard Carpenter, and Karen E. Pollok at Indiana University for their generous contributions in sharing reagents, equipment, and valuable insights during the study.

## Supplementary Information

**Supplementary Fig. S1.**
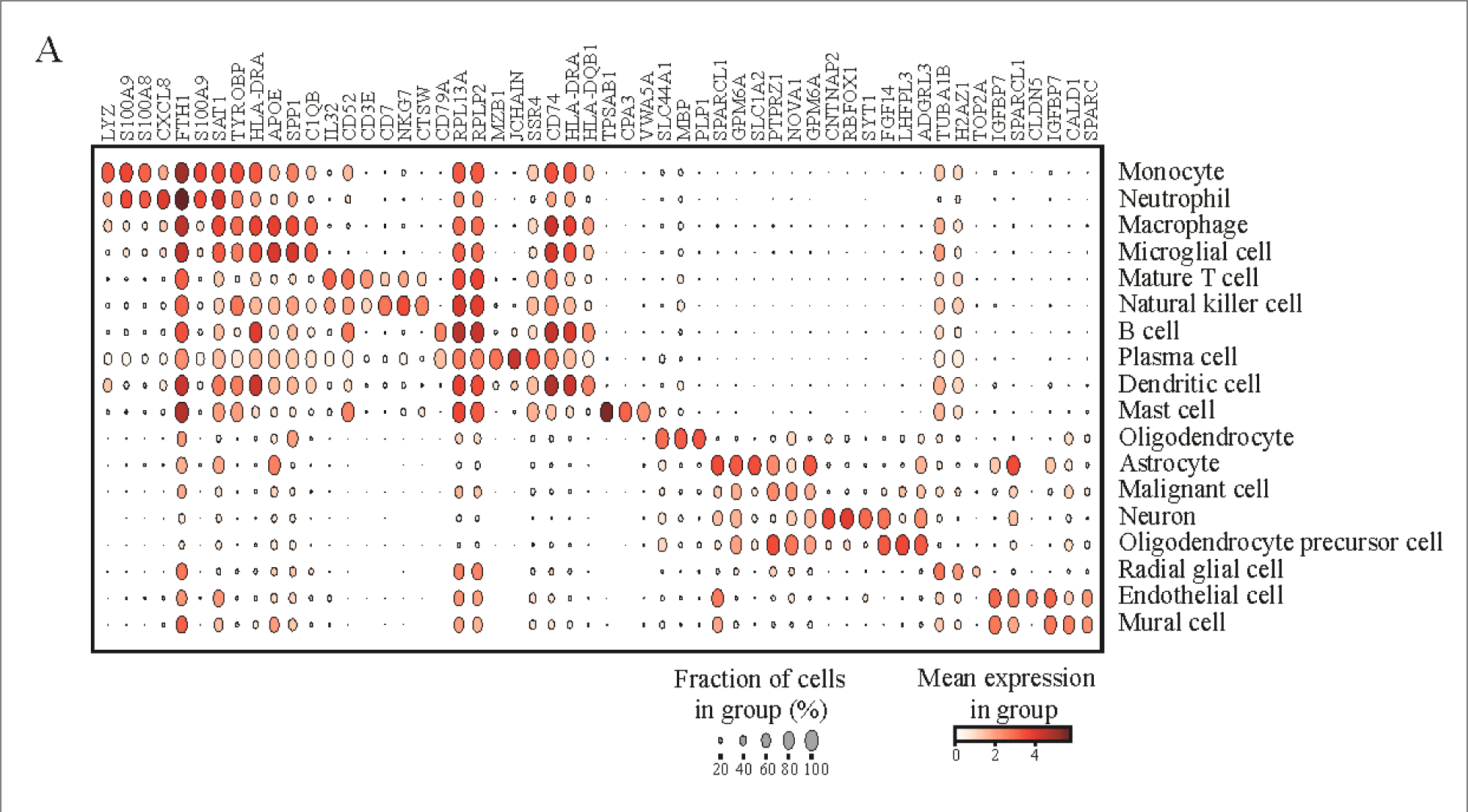
Cell type-specific markers and their expression in GBM. (A) Dot plot showing the expression of cell type-specific markers across different cell populations identified in the single-cell RNA-seq data. The size of each dot represents the fraction of cells expressing the marker in each cell type, and the color intensity represents the mean expression level within the group.

**Supplementary Fig. S2.**
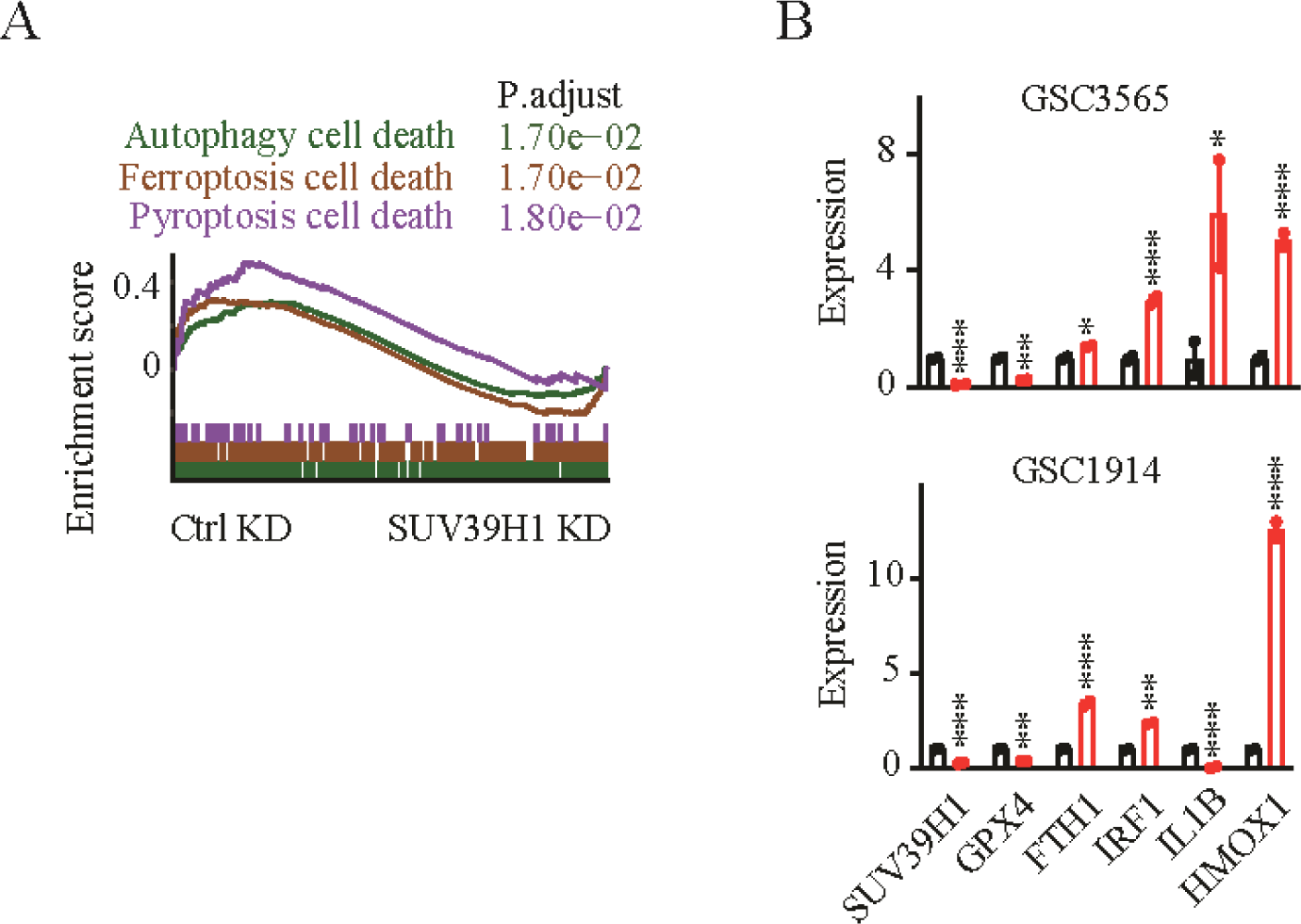
Cell death pathways regulated by SUV39H1 in GSCs. (A) GSEA plot showing enrichment of cell death pathways in SUV39H1 KD GSCs. (B) qPCR detection of cell death-related genes in GSC3565 and GSC1914 cells. *P < 0.05, **P < 0.01, ***P < 0.001.

**Supplementary Table S1. Information for 4 normal and 9 GBM tissues.**

**Supplementary Table S2. qPCR primers.**

**Supplementary Table S3. List of markers used for cell type annotations.**

